# Denaturants Alter the Flux through Multiple Pathways in the Folding of PDZ Domain

**DOI:** 10.1101/221945

**Authors:** Zhenxing Liu, D. Thirumalai

## Abstract

Although we understand many aspects of how small proteins (number of residues less than about hundred) fold, it is a major challenge to understand how large proteins self-assemble. To partially overcome this challenge, we performed simulations using the Self-Organized Polymer model with Side Chains (SOP-SC) in guanidinium chloride (GdmCl), using the Molecular Transfer Model (MTM), to describe the folding of the 110-residue PDZ3 domain. The simulations reproduce the folding thermodynamics accurately including the melting temperature (*T_m_*), the stability of the folded state with respect to the unfolded state. We show that the calculated dependence of ln *k_obs_* (*k_obs_* is the relaxation rate) has the characteristic Chevron shape. The slopes of the Chevron plots are in good agreement with experiments. We show that PDZ3 folds by four major pathways populating two metastable intermediates, in accord with the kinetic partitioning mechanism. The structure of one of the intermediates, populated after polypeptide chain collapse, is structurally similar to an equilibrium intermediate. Surprisingly, the connectivities between the intermediates and hence, the fluxes through the pathways depend on the concentration of GdmCl. The results are used to predict possible outcomes for unfolding of PDZ domain subject to mechanical forces. Our study demonstrates that, irrespective of the size or topology, simulations based on MTM and SOP-SC offer a framework for describing the folding of proteins, mimicking precisely the conditions used in experiments.

## Introduction

The most common way of initiating folding (unfolding) of proteins in ensemble and single molecule experiments is by decreasing (increasing) the concentration of denaturants. Thus, direct comparison with experiments is only possible if simulations are done using models that take the effects of denaturants into account.^1^ Although atomic detailed simulations hold the promise of quantitative description of denaturant-induced folding or unfolding,^2-6^ currently the only available method for obtaining the thermodynamics and folding kinetics of proteins, even for proteins as large as GFP,^7^ is the Molecular Transfer Model (MTM) in combination with coarse-grained SOP-SC representation of polypeptide chain.^8,9^ The theoretical basis for the success of the MTM has been explained elsewhere.^10^ Applications of MTM to probe folding of a variety of proteins have yielded quantitative agreement with experiments ^7,8,11-13^ which attests to the efficacy of the MTM.

One of the early applications of MTM was the demonstration that the Chevron plot of the 56-residue srcSH3 domain could be reproduced nearly quantitatively.^9^ However, the extension of these calculations to proteins with more than hundred residues has been difficult even using the simplified coarse-grained SOP-SC models. One of the goals of this study is to overcome this challenge. The second problem that we address is to establish the folding mechanism as a function of denaturant concentration for a large single domain protein. If the folding mechanism involve parallel pathways, as theoretical and computational studies have firmly established,^14-17^ then are the fluxes through the pathways modulated by changing the external conditions such as denaturant concentration or mechanical forces? Understanding the origin of parallel folding and unfolding pathways, and how they are altered by environmental changes, is important in establishing the generality of the protein folding mechanisms. Here, we investigate the denaturant-dependent folding and unfolding of PDZ3, a protein with110 residues.

PDZ domains, found in many cell junction-associated proteins, are a large family of globular proteins that mediate protein-protein interactions and play an important role in molecular recogniton.^18-21^ The folding of PDZ3 domain, a member of this family, has been studied both by experiments and computations. ^22-26^ The constructs used in these experiments differ. For example, Bai and coworkers^22^ used the construct with two additional β-strands at the C terminal, which are not found in the native PDZ3 domain. Two recent experiments^27,28^ have shown that much of the folding properties, such as the existence of an intermediate or the nature of the transition states, are not greatly affected in the presence or absence of non-native structural elements. In GmdCl-induced equilibrium unfolding experiments of PDZ3 or its variants, the folding transition appears to be highly cooperative, seemingly displaying a simple two-state behavior. In the Chevron plot of PDZ3 domain in GdmCl solution at pH= 6.3, both the folding and unfolding arms are linear functions of GdmCl concentrations [*C*], indicating no detectable intermediate states at the ensemble level. However, native-state hydrogen exchange experiments reveal hidden intermediate states under native conditions.^22^ Interestingly, the addition of potassium formate at pH= 2.85 induces a rollover in the unfolding arm in the Chevron plot, suggestive of an intermediate.^24,25^ This finding is reminiscent of the salt-induced detour found in the folding of the protein S6. ^29^ In the presence of potassium phosphate at pH= 7.5, there are two thermal unfolding transitions in DSC experiments, further demonstrating the existence of intermediate states, at least in the presence of salt.^26^ Based on these experiments, we surmise that the PDZ3 domain folding must occur by multiple pathways, even if they are hard to detect in generic ensemble experiments. If there are multiple pathways, is it possible that the fluxes through these pathways could change by altering the concentration of denaturant in this large single domain as reported for a two-domain protein^30^? Here, we answer this question in the affirmative using MTM simulations in the folding of PDZ3 domain as a function of GdmCl concentration, after establishing that the SOP-SC simulations capture the key findings in ensemble experiments.

We combine simulations of coarse-grained off-lattice SOP-SC model^31-33^ and the molecular transfer model^8-10^ to decipher the folding mechanism of PDZ3 domain. The calculated fractions of molecules in the native basin of attraction(NBA), *f_NBA_*, and the unfolded basin of attraction(UBA), *f_UBA_* as a function of the denaturant concentration [*C*] are in excellent agreement with experiments. In addition, we find that a small fraction, *fI*_BA_, of an intermediate, not resolved in ensemble experiments, is populated in equilibrium The Tanford β parameters for the two transition state ensembles obtained in simulations are in quantitative agreement with those inferred from experiments.^24,25^ Chevron plot calculated for the first time for a protein with over 100 amino acids, allows us to extract the locations of the transition state ensembles (TSEs) in terms of the Tanford β parameters. The calculated free energy profiles suggest that the at low (high) [GdmCl] a less (more) structured TSE is rate limiting. Folding trajectories both in aqueous and in denaturant solutions demonstrate directly the existence of the thermodynamic intermediate states as well as kinetic intermediate states. Our simulations vividly illustrate four parallel folding pathways in molecular detail. We show that the fluxes between the assembly pathways can be modulated by varying the denaturant concentration. Many of our predictions are amenable to experimental test.

## Methods

### SOP-SC model

Our simulations were carried out using the SOP-SC (Self-Organized Polymer-Side Chain) model for the protein.^9,10^ Each residue is represented by two interaction centers, with one centered at the C_α_ position, and the other located at the center of mass of the side chain. The energy function of a conformation in the SOP-SC representation of the polypeptide chain is,

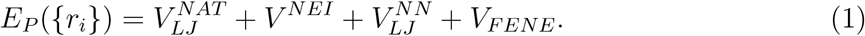

The detailed functional form and the values of the parameters are described elsewhere.^9^

### Molecular Transfer Model

In the MTM the effective free energy function for a protein in aqueous denaturant solution is

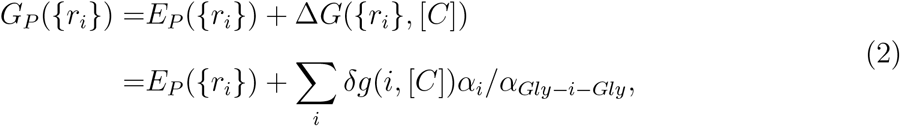

where *Δ_G_*({*r_i_*}, [*C*]) is the free energy of transferring a given protein conformation from water to aqueous denaturant solution, the sum is over all the interaction centers (*i*), *δ_g_*(*i*, [*C*]) is the transfer free energy of interaction center *i*, *α_i_* is the solvent accessible surface area (SASA), and *α_Gly-i-Gly_* is the SASA of the *i^th^* interaction center in the tripeptide *Gly – i – Gly*.

We used the procedure described in detail previously^9,10^ to calculate the thermodynamic properties of proteins in the presence of denaturants.

### Langevin Dynamics

We assume that the dynamics of the protein is governed by the Langevin equation,

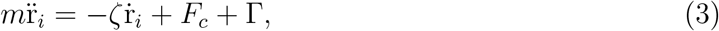

where *m* is the mass of a bead, *ζ* is the friction coefficient, *F_c_* = –*∂E_P_*({*r_i_*})/*∂r_i_* is the conformational force calculated using Eq. (1), Γ is the random force with a white noise spectrum. To enhance sampling, we used Replica-Exchange Molecular Dynamics (REMD)^34-36^ to carry out thermodynamics sampling at low friction coefficient *ζ_L_* = 0.05m/*τ_L_*.^37^ Here, *τ_L_* is the unit of time (see below) for use in the computation of thermodynamic quantities, and *m* is the average mass of the beads. It is obvious that the precise values of these two quantities do not play a role in the determination of the equilibrium properties of interest. In the underdamped limit, we employ the Verlet leap-frog algorithm to integrate the equations of motion.

### Brownian Dynamics

To obtain a realistic description of the kinetics of folding or unfolding, we set *ζ_H_* = 50*m/τ_L_*, which approximately corresponds to the value of the friction coefficient in water. ^38^ At the high *ζ* value where the inertial forces are negligible, we use the Brownian dynamics algorithm^39^ to integrate equations of motion using

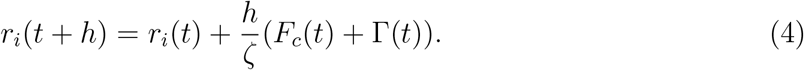

It is important to note for obtaining kinetic properties the systematic force, *F_c_* in Eq. 4 is,

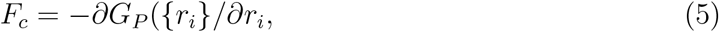

where *G_P_* ({*r_i_*}) is given in Eq. (2).

### Time scales

In the high *ζ* limit the unit of time is 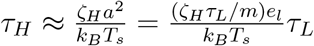. Following Veitshans,^38^ we chose *e_l_* = 1 *kcal/mol*, average mass *m* =1.8 × 10^−22^ *g, a* = 4 Å, which makes *τ_L_* = 2 *ps*. For *ζ_H_* = 50 *m/τ_L_*, we obtain *τ_H_* = 164 *ps*. These estimates are used to obtain estimates of the folding times from our Brownian dynamics simulations.

In Langevin dynamics simulations, the integration time step, *h* = 0.05*τ_L_*, whereas in the Brownian dynamics simulations, *h* = 0.1*τ_H_* (Eq. (4)).

## Results

The structure of the *N* = 110 residue PDZ3 domain from PSD-95 is shown in Fig-1A (PDB ID 1BFE). In contrast to the typical structure of a PDZ domain, it has one additional helix (*α*_3_ and a two-stranded *β* sheet formed between two short strands, *β*_7_ and *β*_8_ at the C terminal. They are colored in red in Fig-1A.

**Fig-1:**
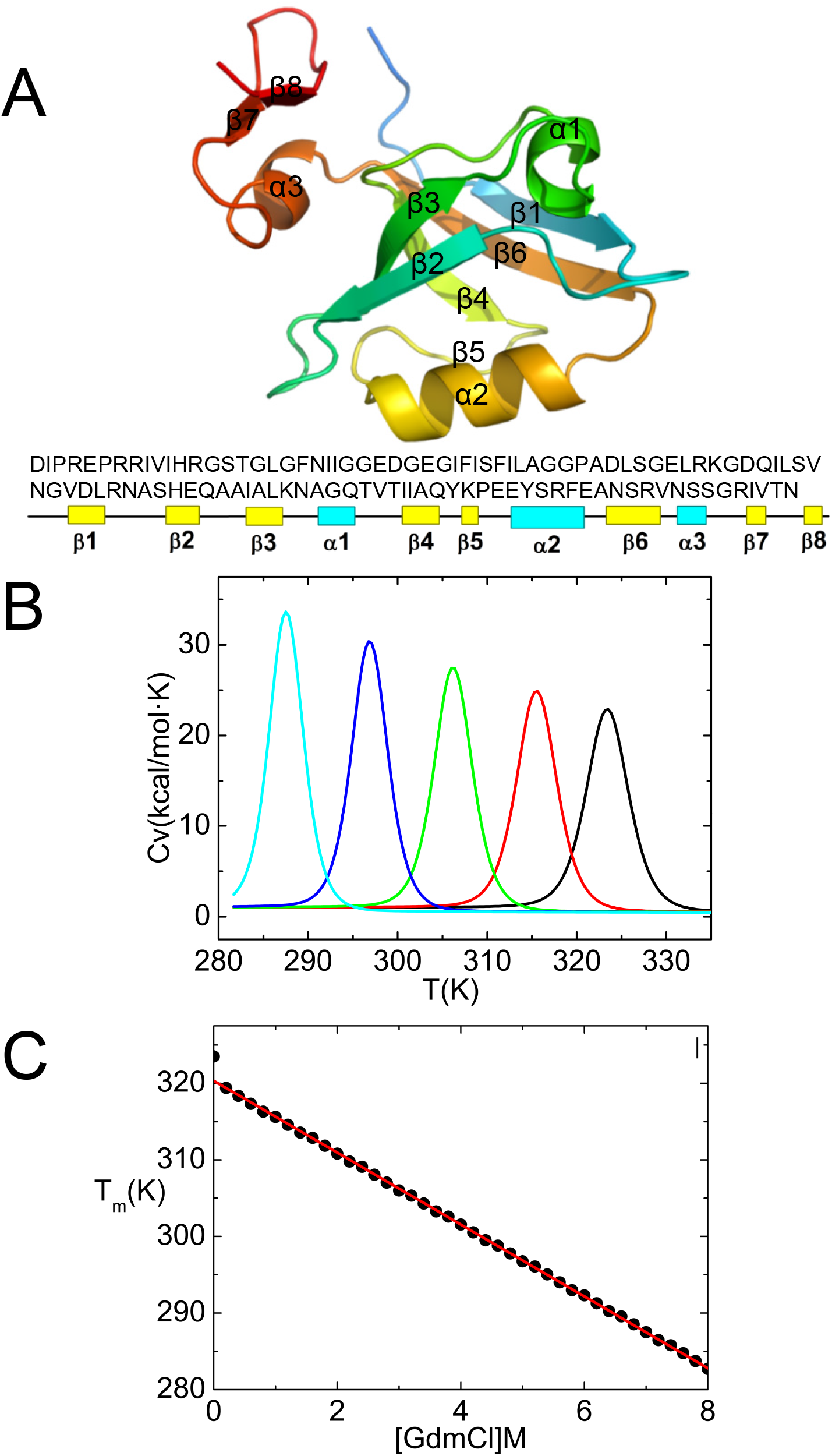
(A) Cartoon representation of PDZ3 domain (PDB code: 1BFE). The sequence used in the simulations is given below using a one letter code for amino acids. (B) Heat capacity for different values of the GdmCl concentration, [*C*]. The values of [*C*] measured in M from right to left are 0, 1, 3, 5 and 7. (C) The dependence of the melting temperatures, T_m_[*C*]s on [*C*].

### Melting temperatures

The melting temperature, identified with the peak in the heat capacity *C_v_*(*T*) at [*C*] = 0 (black line in Fig-1B) is *T_m_* = 323*K*, which is in reasonable agreement with the experimentally measured *T_m_* = 344K.^26^ Similarly, by associating the melting temperatures with the peaks in the heat capacity at different values of [*C*] (Fig-1 B) we determined the dependence of *T_m_[C*] on [*C*]. It is clear from Fig-1 C that *T_m_*[*C*] is linearly dependent on [*C*] (see figure caption for the parameters of the fit).

### Boundaries between the distinct thermodynamic states

To define the Native Basin of Attraction (NBA) and the basins corresponding to the unfolding (UBA) and intermediate states (IBA), we obtained the free energy profile *G*(*χ*) as a function of the structural overlap function, *χ*, which serves as an order parameter. The structural overlap function 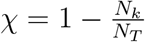 where,

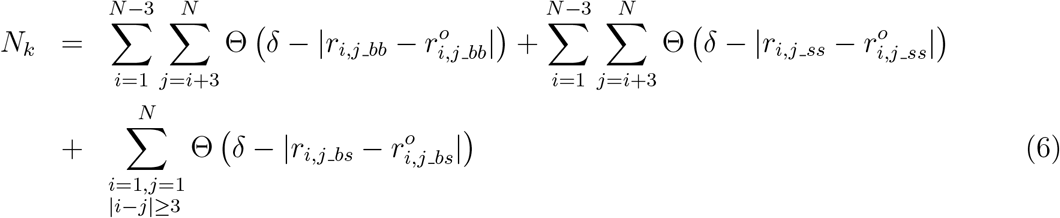

In Eq. (6), Θ(*x*) is the Heavyside function. If 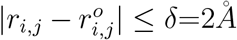, there is a contact. *N_k_* is the number of contacts in the *k^th^* conformation and *N_T_* is the total number in the folded state. The profile *G*(*χ*) at *T_m_*[0] (Fig-2A) shows that the conformations can be classified into three groups separated by the black vertical lines; 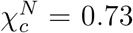 and 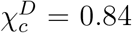 are the cutoff values separating the NBA, UBA, and IBA. If 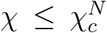, the corresponding conformation belongs to the NBA and if 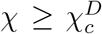, the conformation belongs to UBA, and all the other conformations are grouped into IBA.

**Fig-2:**
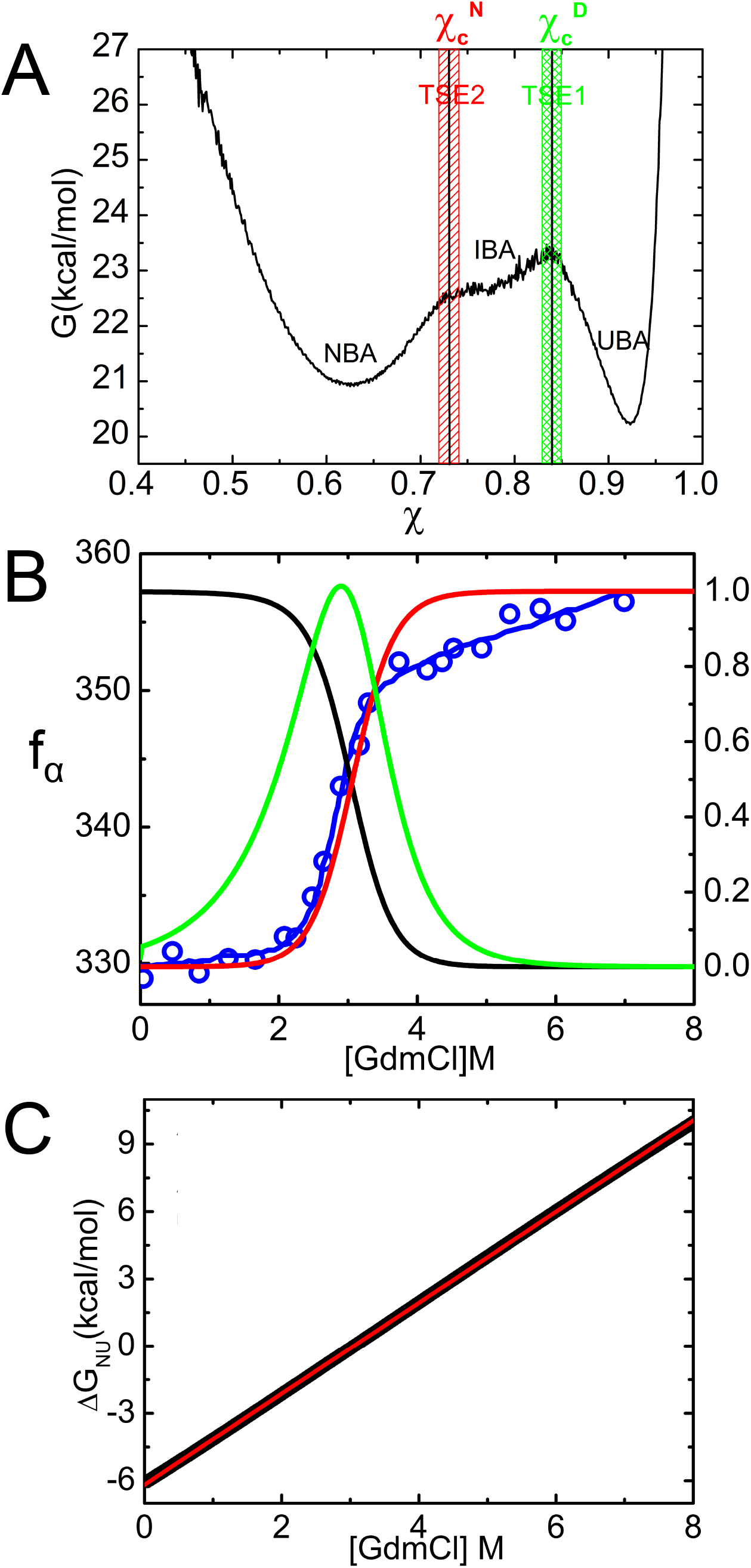
(A) Free energy profile at the melting temperature *T_m_*, as a function of the χ, structural oder parameter. 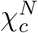 and 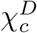 identify the boundaries between the states. The locations of the TSE1 between the NBA and IBA and TSE2 separating the IBA and UBA are labelled. (B) Fraction of molecules in the NBA (black, right scale), UBA (red, right scale) and IBA (green, right scale × 0.025) as a function of GdmCl concentration, [*C*]. For comparison, experimentally monitored maximum wavelength of the fluorescence at different concentrations of GdmCl (blue, left scale) is also shown. Although it is not directly equal to *f_UBA_*, it is correlated to *f_UBA_*. (C) The [*C*] dependence of the free energy of stability of the native state with respect to the unfolded state, Δ*G_NU_*.

### GdmCl dependence of thermodynamic stability and m-value

In order to compare with experiments, we simulated the effects of GdmCl using the Molecular Transfer Model (MTM).^8^ Following our previous studies,^8-10^ we choose a simulation temperature, *T_s_*, at which the calculated free energy of stability of the native state (*N*) with respect to the unfolded state (*U*), Δ*G_NU_*(*T_s_*) (*G_N_*(*T_s_*) – *G_U_*(*T_s_*)) and the measured free energy at *T_E_* (=298K) Δ*G_NU_*(*T_e_*) coincide. For PDZ3, Δ*G_NU_*(*T_E_* = 298K) = –7.4*kcal/mol* at [*C*] = 0,^22^ which results in *T_s_* = 306K, which is close to *T_E_*. It is worth emphasizing that besides the choice of *T*s** no other parameter is adjusted to fit any experimental data.

With *T_s_* = 306*K* fixed, we calculated the dependence of the fraction of molecules in the NBA, *f_NBA_*([*C*],*T_s_*), in the UBA, *f_UBA_*([*C*],*T_s_*), and in the IBA, *f_IBA_*([*C*],*T_s_*), on [*C*] (Fig-2B). The midpoint concentration, *C_m_*, obtained using *f_UBA_*([*C_m_*], *T_s_*) = 0.5 is [*C*]=3.05M, which is very close to 2.90M, measured in experiments.^22^ For comparison, the experimentally monitored maximum wavelength of the fluorescence at different concentrations of GdmCl is also shown (blue, left scale in Fig.2B). Although it is not a direct measure of *f_UBA_*, it is correlated to *f_UBA_*, which in turn is in good agreement with the result based on simulations. The ability to reproduce reasonably accurately experimental measurements further establishes the efficacy of the MTM and SOP-SC simulations in capturing the folding thermodynamics of single domain proteins in general, and PDZ3 in particular.

The small value of *f_IBA_*([*C*],*T_s_*), compared to *f_NBA_*([*C*],*T_s_*) and *f_UBA_*([*C*],*T_s_*), explains why the intermediate state is hard to detect in the equilibrium denaturation experiments. ^22^ Our finding that *f_IBA_*([*C*],*T_s_*) is small is consistent with the observed protein-concentration dependent thermal unfolding in DSC experiments, where at low PDZ3 concentration only one transition is observed as the associated intermediate is not substantially populated. ^26^

The native state stability with respect to *U*, Δ*G_NU_*([*C*])(= *G_N_*([*C*]) – *G_U_*([*C*])), is calculated using 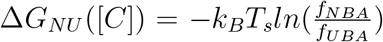. The linear fit, Δ*G_NU_*([*C*]) = Δ*G_NU_*(0) + *m*[*C*], yields Δ*G_NU_*([0]) = –6.18*kcal/mol* and *m* = 2.03*kcal/mol · M* (Fig. 2C), which is in reasonable agreement with experimental estimate *m* = 2.50*kcal/mol · M*.^22^ In light of a recent experiment showing that the truncation of the *α_3_* helix only modestly destabilizes the native state,^27^ we surmise that the addition of extra two *β* strands at the C terminal probably does not significantly affect the stability of PDZ3, and thus the value of Δ*G_NU_*([0]).

### Free energy profiles as function of the order parameter *χ*

To illustrate how GdmCl changes the free energy landscape, we plotted the free energy profiles as functions of *χ* at different [*C*] at *T_m_[C*] in Fig-3A and at a fixed temperature, *T_s_* = 306*K* in Fig-3B. We use *χ* in Eq. (6), the microscopic order parameter of the protein, to distinguish between the native, the unfolded and high-energy intermediate states.^40^ Fig-3B shows that, at low [GdmCl], the first transition state ensemble, TSE1, is rate limiting for folding. However, at high [GmdCl] the second transition state ensemble, TSE2, is rate limiting. These findings agree qualitatively with the energy diagram for the folding reactions proposed elsewhere. ^24^ The movement of the transition states with changing concentration is in accord with the Hammond postulate.

**Fig-3:**
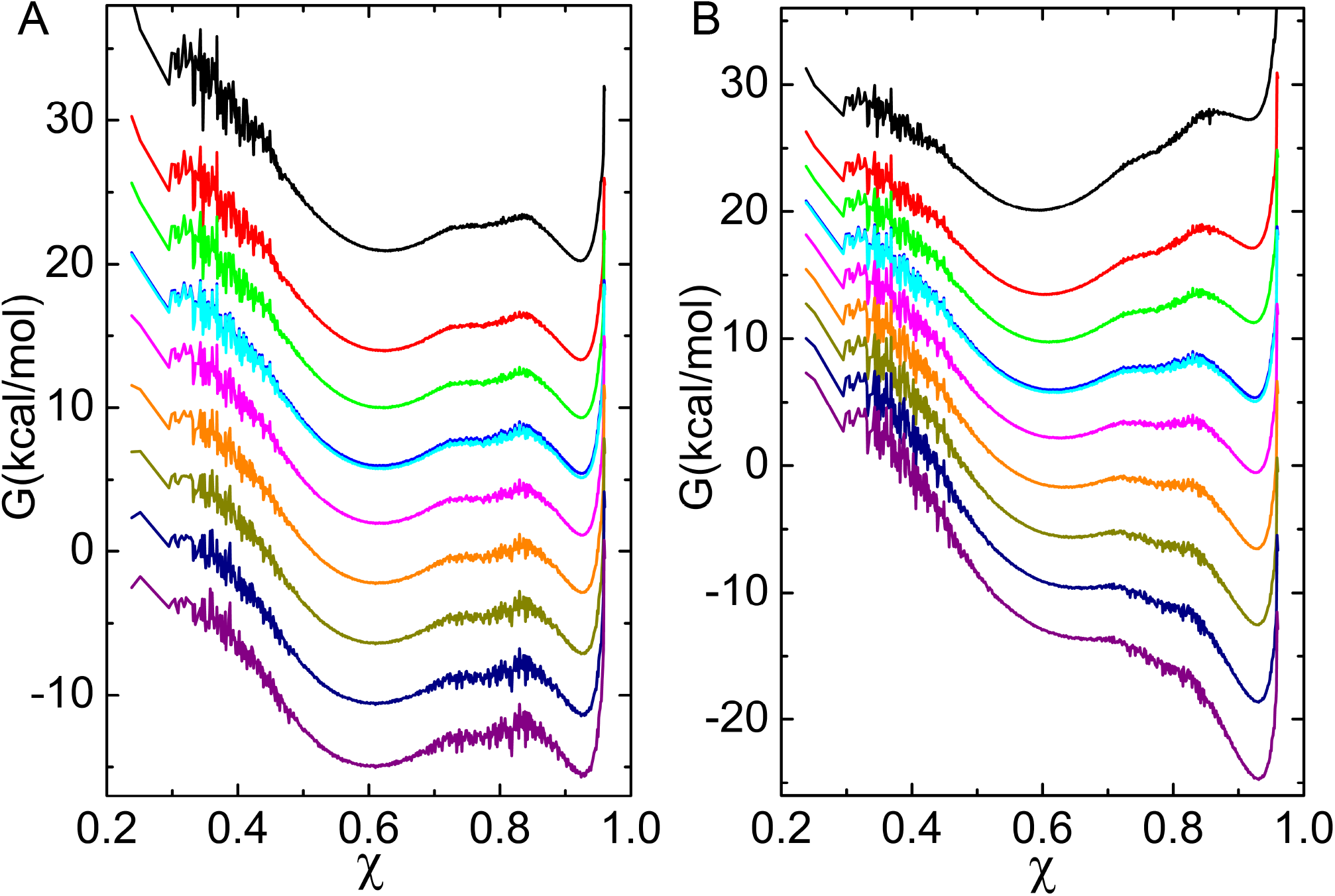
Free energy profiles as functions of *χ* at different [*C*]. (A) *T* = *T_m_*. (B) *T* = *T_s_*. The values of [*C*] measured in molar units from top to bottom are 0, 1, 2, 3, 3.05, 4, 5, 6, 7, 8.

### Structures of the transition state ensembles (TSEs)

The free energy profile as a function of *χ* in Fig. 2A, suggests that there are two barriers. The ensembles of conformations at their locations, grouped as TSE1 and TSE2, are shown by the shaded areas. The global characteristic of the TSE in ensemble experiments is usually described using the Tanford parameter, *β*. From the observed chevron plot, 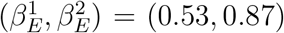^24^ or (0.56, 0.90) for TSE1, TSE2 respectively.^25^ It is generally assumed, that *β* is related to the buried solvent accessible surface area (SASA) in the TSE. For the TSE obtained in our simulations, we calculated the distribution *P*(Δ_*R*_) (Fig-4A), where Δ_*R*_ = (Δ_*U*_ – Δ_*TSE*_)/(Δ_*U*_ – Δ_*N*_) with Δ_*U*_, Δ_*TSE*_, *Δ_*N*_* are the SASA in the DSE ([*C*] = 8.0M), TSE, and the NBA ([*C*] = 0.0M), respectively. We found that the average 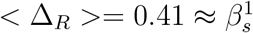 for TSE1 and 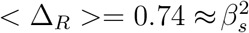 for TSE2, which are in qualititative agreement with the experimentally measured values. The small deviations between simulations and experiments may be due to the following reasons. (1) The PDZ3 domain in the simulations has one additional helix and one *β* sheet, which is absent in the construct used in the experiments. These extra structural elements are highly flexible even under native conditions (upper left in Fig-4B), which lowers < Δ_R_ >. (2) When fitting the chevron plot to obtain *β* in experiments, both 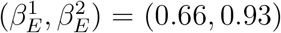 and (0.3, 0.8) give reasonable fits, indicating a range of *β* can describe the experimental data.^24^ Given these observations, we surmise that the Tanford *β* calculated using the simulation data is in reasonable agreement with experimental values.

**Fig-4:**
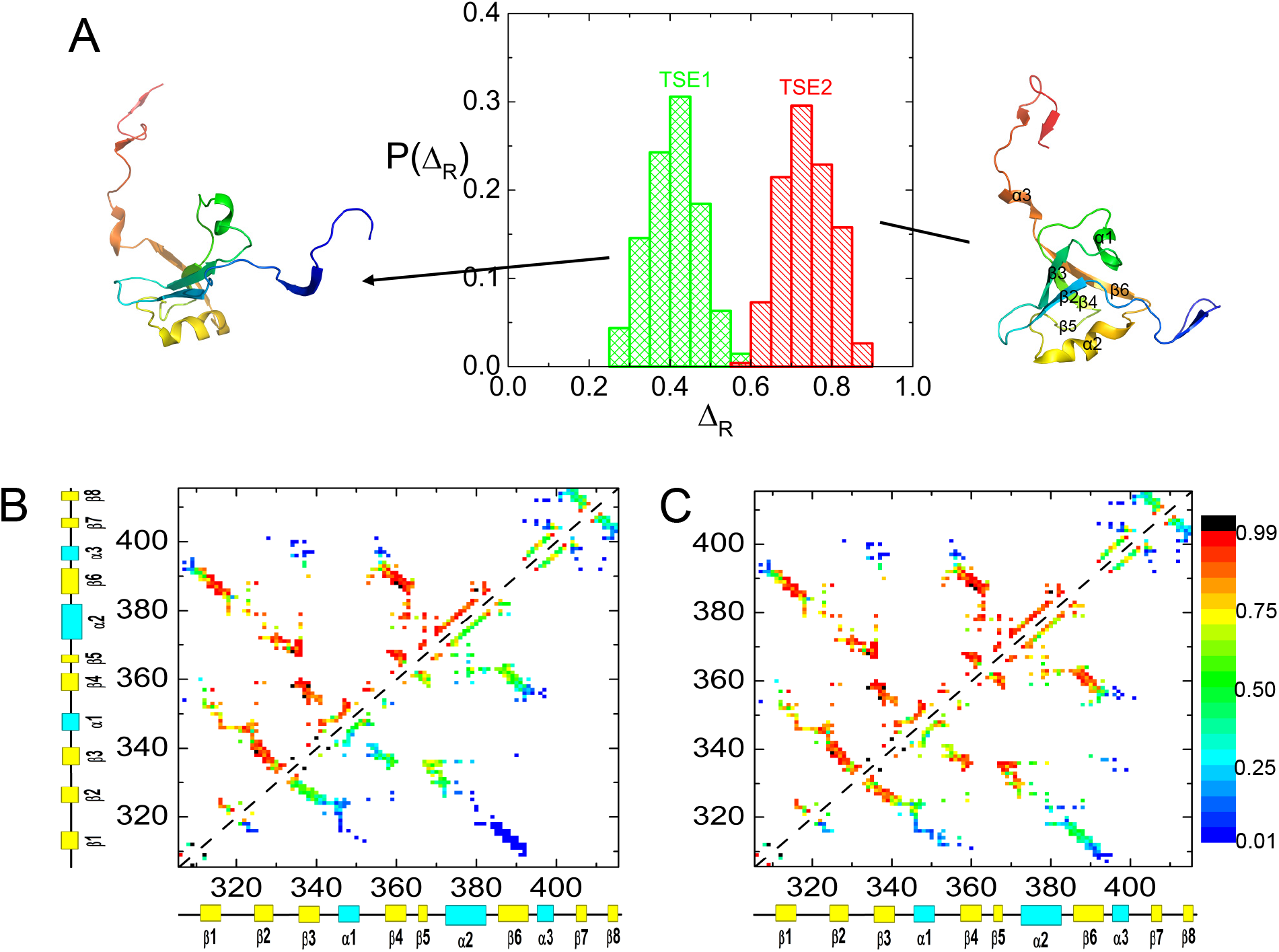
Transition state ensembles. (A) Distribution *P*(Δ_*R*_) of the Δ_*R*_ = (Δ_*U*_ – Δ_*TSE*_)/(Δ_*U*_ – Δ_*N*_), which is the fraction of buried solvent accessible surface area relative to the unfoled structures. The average < Δ_*R*_ >= 0.41 for TSE1 and < Δ_*R*_ >= 0.74 for TSE2. Snapshot of TSE1(TSE2) is shown on the left(right). (B) Contact maps of the native state ensemble (upper left) and the one for the TSE1 (lower right). (C) The same as B except for the TSE2.

Fig-4B and Fig-4C show the contact maps obtained from the TSE1 and TSE2, respectively. It is clear that, relative to the native state (upper left) the TSE1 structures, populated at low [*C*], is moderately structured. In contrast, the structures are ordered to a greater extent in the TSE2. The contacts between *β*1 – *β*6 have very low probabilities of formation indicated by the major blue region in Fig-4B, and have moderate formation probabilities indicated by the major green region in Fig-4C. One representative structure for TSE1 is shown on the left of Fig-4A, where we can see that *β*_2_, *β*_3_, *β*_4_, *β*_5_, *β*_6_, *α*_1_, *α*_2_ and *α*_3_ are packed loosely with *β*_1_, *β*_7_, *β*_8_ forming no contact with the core of the protein. Note that *β*_7_, *β*_8_ and *α*_3_ are the extra regions in our simulations compared to the typical structure of PDZ domain. A representative structure for TSE2 is shown on the right of Fig-4A, where the core of the native topology is well established except for *β*_1_, *β*_7_ and *β*_8_. We should point out a minor discrepancy between our results and a previous study. ^25^ We find that *β*1 is unstructured in the TSE2 but is found to be structured by Jemth.^25^

### Folding kinetics and the Chevron Plot

We calculated the [*C*]-dependent folding (unfolding) rates from folding (unfolding) trajectories, which were generated from Brownian dynamics using the effective energy function, *G_P_*({*r_i_*}) (see Methods). From sixty (one hundred for [*C*] = 0) folding trajectories, the fraction of unfolded molecules at time *t*, is computed using 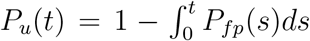, where *P_fp_*(*s*) is the distribution of first passage times. We fit *P_u_*(*t*) ≈ *e*^−*tk_f_*^under folding conditions ([*C*] < *C_m_*), from which *k_f_*[*C*] can be extracted. Similarly, a single exponential fit for unfolding conditions ([*C*] > *C_m_*) yields *k_u_*[*C*]. At high (low) [*C*], we can approximate *k_obs_* = *k_f_*([*C*]) + *k_u_*([*C*]) as *k_u_*([*C*]) (*k_f_*([*C*])). We globally fit the relaxation rate, *k_obs_* using ln*k_obs_*=ln[*k_f_*([0])*e^−m_f_/RT^* + *k_u_*([0])*e*^−*m_u_*[*C*]/*RT*^], where *m_f_* (*m_u_*) is the slope of the folding (unfolding) arm with ln*k_f_*=ln*k_f_*(0) – *m_f_*[*C*]/RT and ln*k_u_*=ln*k_u_*(0) + *m_u_*[*C*]/*RT*.

A plot of ln *k_obs_* as a function of [*C*] over a wide concentration range (0M ≤ [*C*] ≤ 8.0M) shows a classic Chevron shape (Fig-5) observed in several experiments for a number of proteins. In the range [*C*] ≤ 1.5M, *k_u_* ≪ *k_f_*, so that *k_obs_* ≈ *k_f_* and similarly for [*C*] above 4.5M, *k_obs_* ≈ *k_u_*. In the transition region (2.0M ≤ [*C*] ≤ 4.0M), the folding and unfolding rates are too small to be reliably calculated even using the SOP-SC simulations. Because the size of the PDZ3 domain is relatively large (110 amino acids) we could not generate folding and unfolding rates reliably around the midpoint even using the SOP-SC model. Comparison of the simulation and experimental results shows that the slopes (from the folding and unfolding arms) of the simulated chevron plot are qualitatively similar to the experimental values.

**Fig-5:**
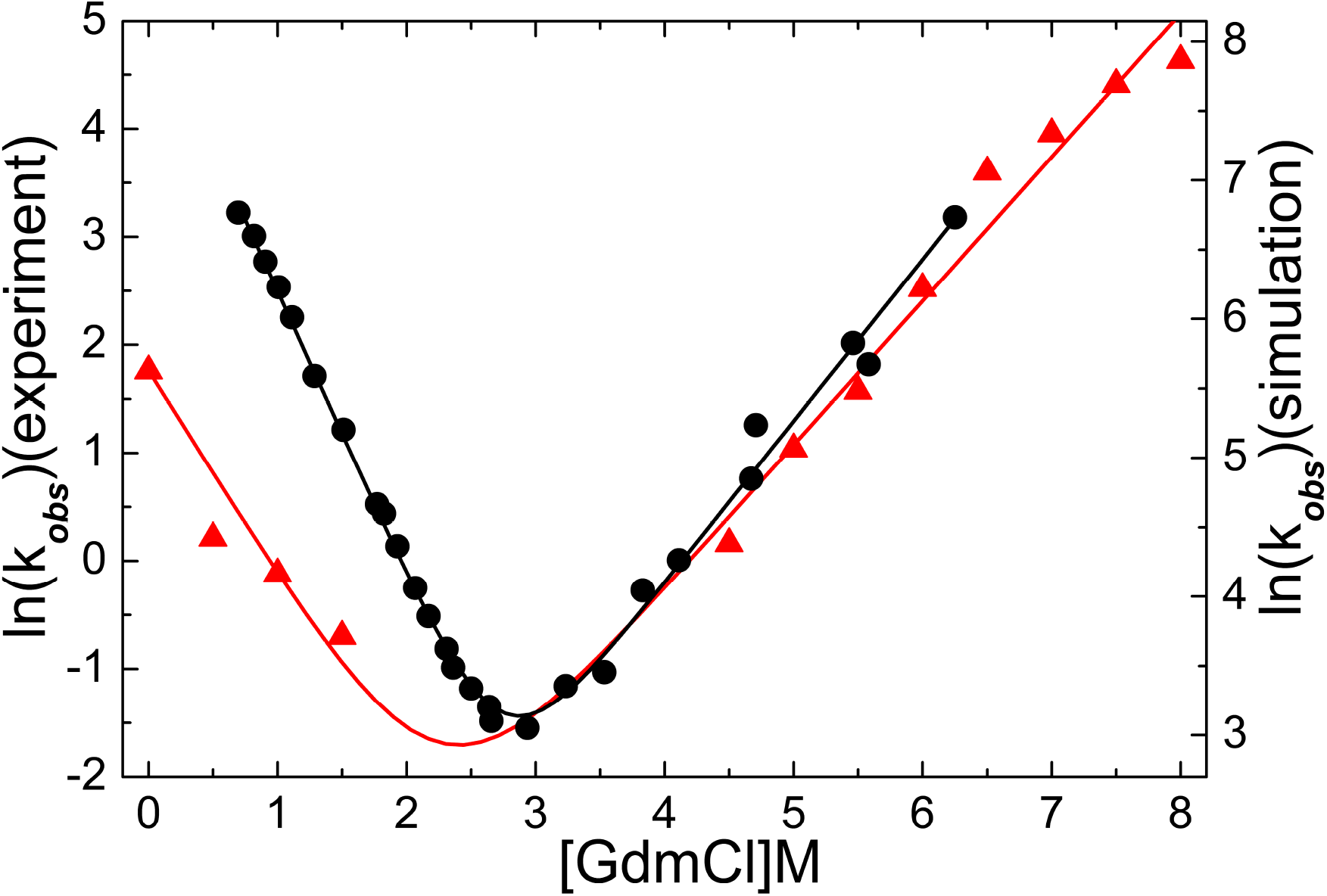
Comparison of chevron plots obtained from simulations and experiment. The scale for the experimental plot (black dots and line) for *lnk_obs_* is on the left, and for the simulation (red triangles and line) is on the right.

From the slope of the folding arm (simulation results in Fig-5), we obtain *m_f_* = 0.91*kcal/mol· M and m_u_* = 0.63*kcal/mol · M* from the unfolding arm. The corresponding experimental values are 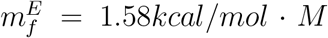 and 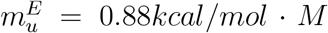.^22^ The agreement between experiments and simulations for the slope of the folding arm is reasonable and the agreement for the unfolding arm slope is fair. Since the the fraction of molecules in IBA is negligibly small, both thermodynamics and kinetics simulations can be approximately described by a two-sate model and hence we expect *m ≈ m_f_* + *m_u_*. From the simulated Chevron plot, we obtain *m* ≈ 1.54*kcal/mol · M*, which differs by ~ 24% from *m* = 2.03*kcal/mol · M* obtained from equilibrium Δ*G_NU_*[*C*] calculations (Fig. 2C). In contrast, the relation *m* = *m_f_* + *m_u_* = 2.46*kcal/mol · M*, which is close to *m* = 2.50*kcal/mol · M* found in equilibrium titration experiments. We conclude that our simulations capture only the qualitative features of the denaturant-dependent folding kinetics of PDZ3 domain.

Although MTM simulations reproduce the Chevron shape well, the dependence of *lnk_obs_* on [*C*] does not agree quantitatively with experiments. For instance, *k_f_* [0] from simulations is 278 *s*^−1^, which is only ≈ 1.6 times larger than the extrapolated value 170 *s*^−1^ for [*C*] = 0 from experiment. However, the unfolding rate at [*C*] = 0, *k_u_*[0] from simulations is 0.91 *s*^−1^ while *k_u_* [0] from experiment is 0.0022 *s*^−1^. The simulations overestimate the unfolding rate by about 414 fold compared to experiments, even though the values of the slopes of the unfolding arms from simulations and experiments are reasonably close. It is not easy to theoretically establish the reasons for the large difference between the predicted and experimental values of the unfolding rate, especially considering that the folding rate is accurate. We expect, on general grounds, that both the folding and unfolding rates should differ from measurements because we use coarse-grained models. In several previous studies^9,10,12^ we had argued that the difference between simulations and experiments could be on one or two orders of magnitude. The effective diffusion in our model is greater than would be the case had the solvent been modeled explicitly, which alas is impossible to do using current atomic detailed simulations. The larger predicted value of *k_u_*[0] compared to experiments suggests that the unfolding energy landscape is rugged, which is not accurately captured by the simulations. Assuming that the actual diffusion upon unfolding 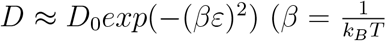 and *ε* is the scale of roughness) the discrepancy between the predicted and experimental *k_u_*[0] implies that the ε ≈ 2.5*k_B_T*. The absence of non-native interactions, which apparently is important for unfolding of PDZ3, could explain the overestimation of *k_u_*[0]. Clearly, the extent of deviation is likely to depend on the protein and the sequence.

### Fluxes through parallel pathways depend on the denaturant concentration

By analyzing the folding trajectories by using *χ* as the progress variable for the folding reaction, we find that PDZ3 folds along four distinct pathways. One representative trajectory for each pathway is shown in Fig-6, where *χ* is displayed as a function of t. In each pathway folding occurs in stages. In addition to the conformations in the UBA and the NBA, we identified two intermediate states (*KIN*_1_ and *KIN*_2_), whose lifetimes vary greatly depending on the pathway. Arrows of each color in Fig-7 represent one folding pathway and the thickness of the arrows represents the probability of the pathway. At [*C*] = 0 (Fig-7A), the dominant pathway P1 is *D* → *KIN*_1_ → *KIN*_2_ → *N* (black arrows), through which ~ 52% of the flux to the native state is channeled. In this pathway, *β* sheets between strands 1, 6, 4 form transiently in *KIN*_1_ state, followed by the consolidation of core *β* sheets between strands 2, 3, 4 and α_2_ in the *KIN*_2_ state. The less probable alternative pathway P2 is *D* → *KIN*_2_ → *N* (red arrows), representing ~ 38% of the trajectories, where folding occurs only through the *KIN*_2_ state. Similarly, in the third probable pathway P3, *D → KIN*_1_ → *N* (green arrows), through which about ~ 10% of the flux to the native state flows, folding occurs only through *KIN*_1_ state.

**Fig-6:**
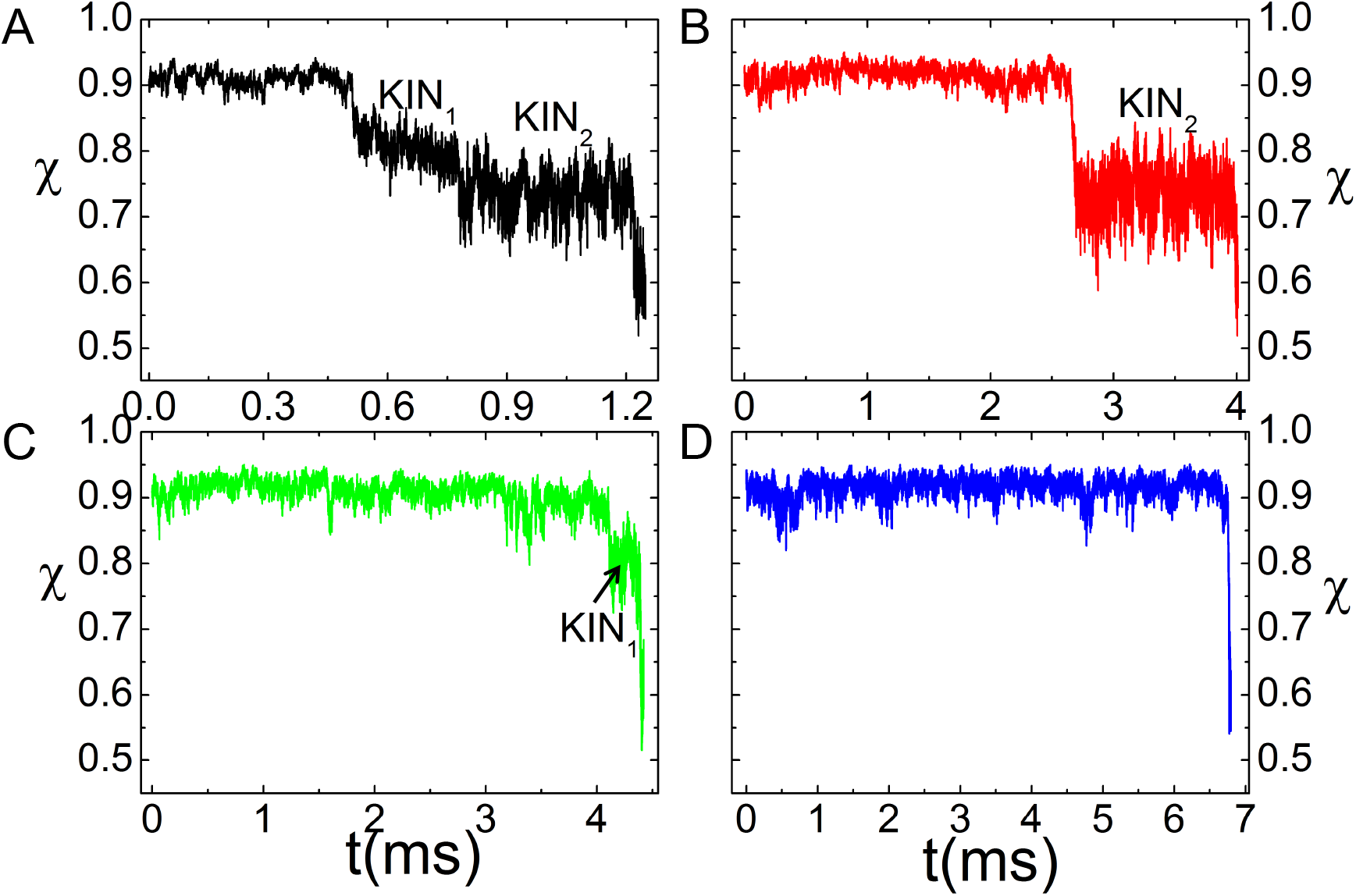
Representative folding trajectories for the four folding pathways. The distinct routes to the folded states are given in terms of the time-dependent changes in *χ*. (A) P1, *D* → *KIN*_1_ → *KIN*_2_ → *N*. (B) P2, *D* → *KIN*_2_ → *N*. (C) P3, *D* → *KIN*_1_ → *N*. (D) P4, *D* → *N*.

**Fig-7:**
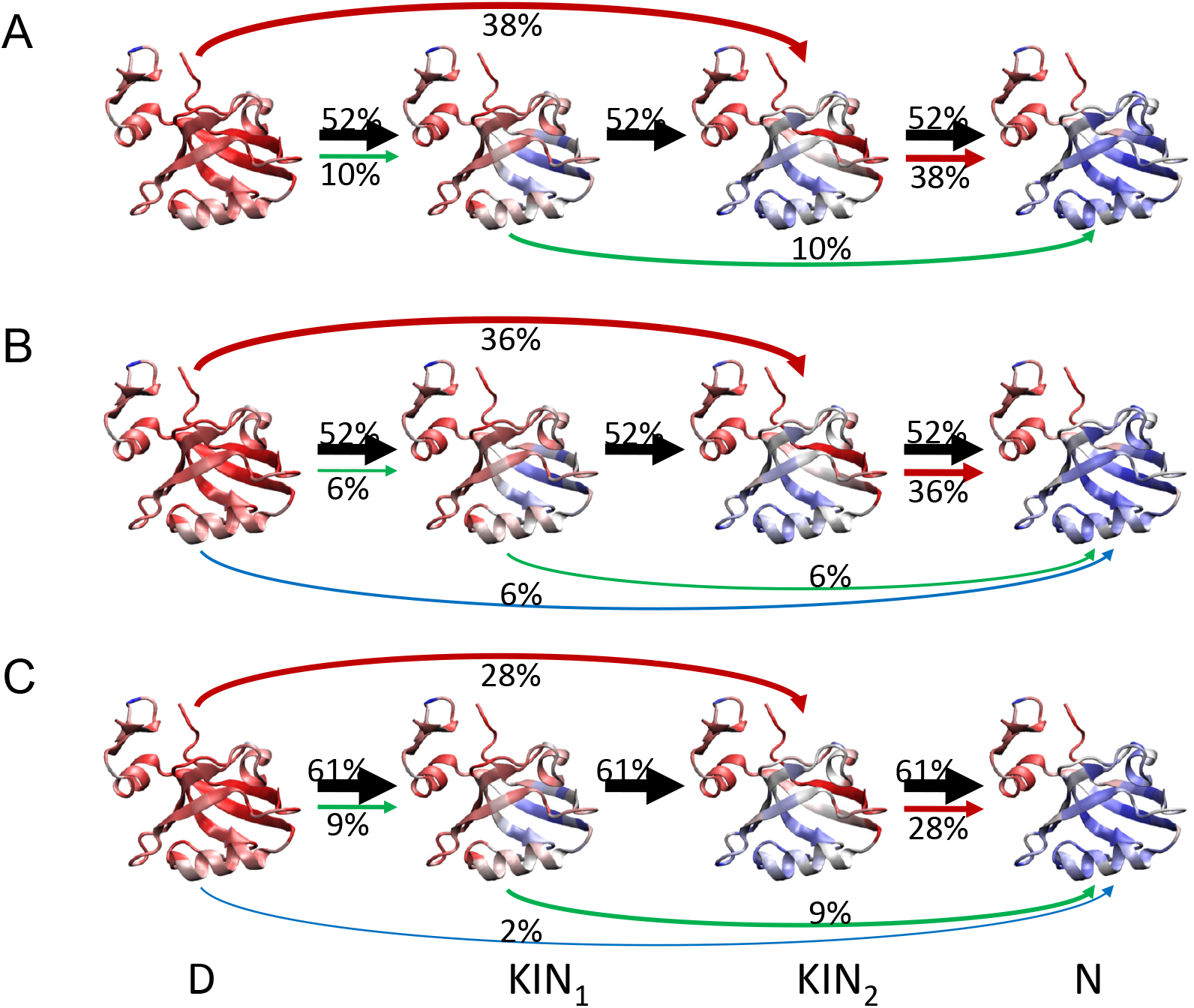
Schematic of the folding pathways and the structural features of the populated states. (A) Folding pathways and fraction of native contacts of all states at [*C*] = 0.0M. Color code: Red, unstructured; Blue: fully structured. The three folding pathways that reach the native state are represented by colored arrows with the their widths representing the flux through the pathways. Different colors represent different folding pathways. (B, C) Same as A but at [*C*] = 0.5M (B) and 1.0M (C) with the states represented by circles.

The flux through P1, identified at [*C*] = 0 remains dominant at [*C*] = 0.5M (Fig-7B, ~ 52%) and [*C*] = 1.0M (Fig-7C, ~ 61%). The P2 and P3 pathways have lesser probabilities ~ 36%, ~ 6% for [*C*]=0.5M and ~ 28%, ~ 9% for [*C*] = 1.0M, respectively. A direct pathway P4, *D* → *N*(blue arrows) is observed with small probabilities (~ 6% for [*C*]=0.5M, ~ 2% for [*C*] = 1.0M). Thus, the PDZ3 domain folds through heterogeneous pathways. Most importantly, the populations of the folding pathways are sensitive to denaturant concentrations. Denaturant-modulated parallel pathways were also observed for adenylate kinase.^30^ Interestingly, such parallel folding has been observed in the folding of both small proteins^41^ and larger proteins. ^30,42,43^

## Discussion

### Post-Collapse kinetic intermediate is structurally similar to an equilibrium intermediate

To illustrate the relationship between the thermodynamically observed intermediate state (*I_EQ_*) and kinetically observed intermediate states (*KIN*_1_, *KIN*_2_), we calculated the average fraction of native contacts at every residue *f_Q_s*. The correlations between the *f_Q_s* for the three states are shown in Fig-8A and Fig-8B. The correlation between *I_EQ_* and *KIN*_1_ is very low (correlation coefficient, R=0.3), indicating that at the early stages of folding a variety of compact but structurally diverse states are explored. The observation that in the initial stages of organization a heterogeneous mixture of states with small thermodynamic states are sampled is consistent with early atomic detailed simulations on cytochrome c. ^44^

**Fig-8:**
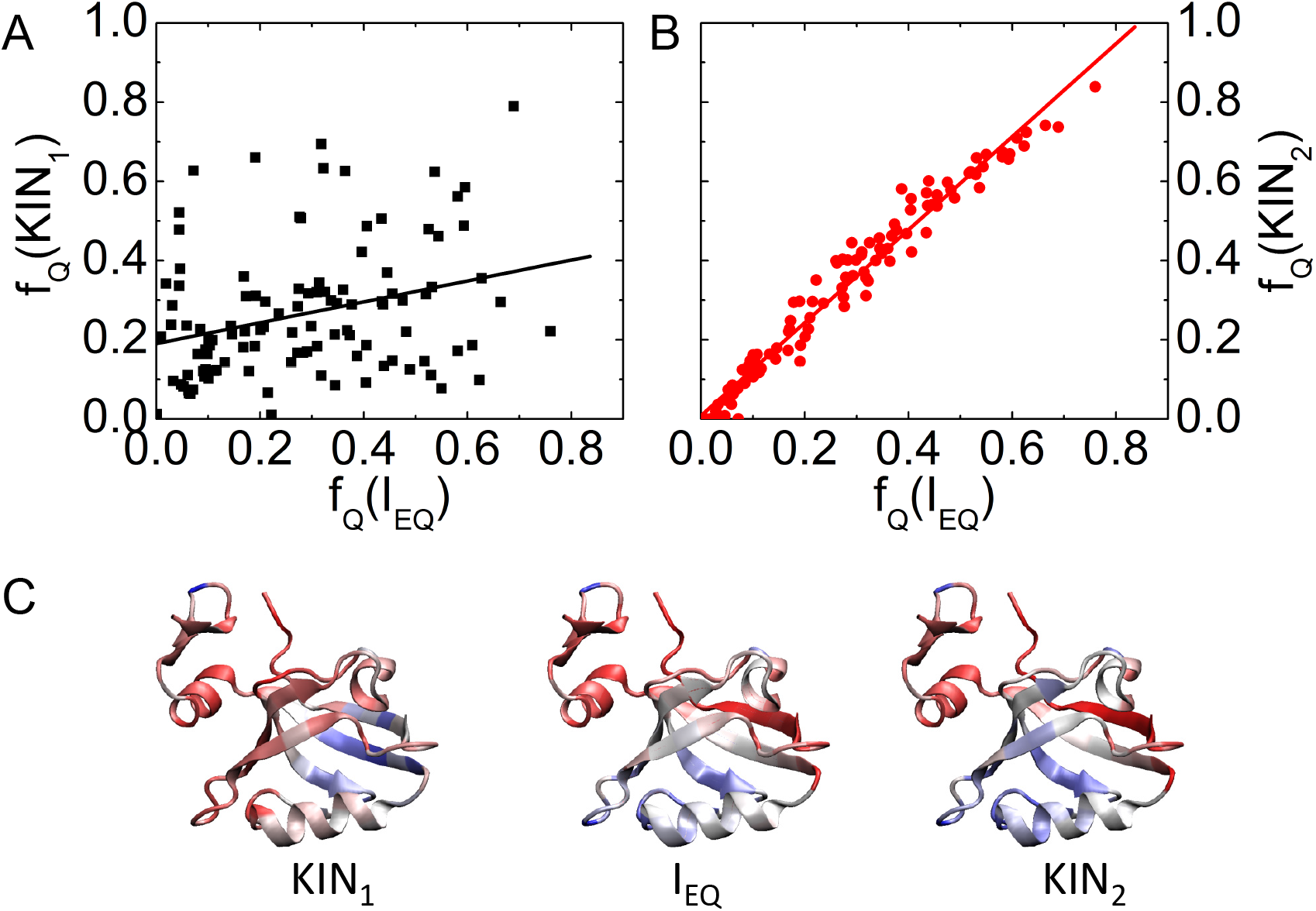
Correlation between equilibrium *I_EQ_* and kinetic intermediates, *KIN*_1_ and *KIN*_2_ using the average residue-resolved fractions of native contacts, *f_Q_s*. (A) Plot of *f_Q_* between *KIN*_1_ and *I_EQ_*. The heterogeneous states populated during the early stages of folding results in little structural correlation between these states.Same as (A) except this plot shows correlation between *KIN*_2_ and *I_EQ_*. At late stages (post collapse) the structures of the intermediates at equilibrium and during kinetics coincide. (C) Sample structures for *KIN*_1_, *I_EQ_*, and *KIN*_2_ are displayed. Red represents structures not present in the native state and blue is native-like.

**Figure 9:**
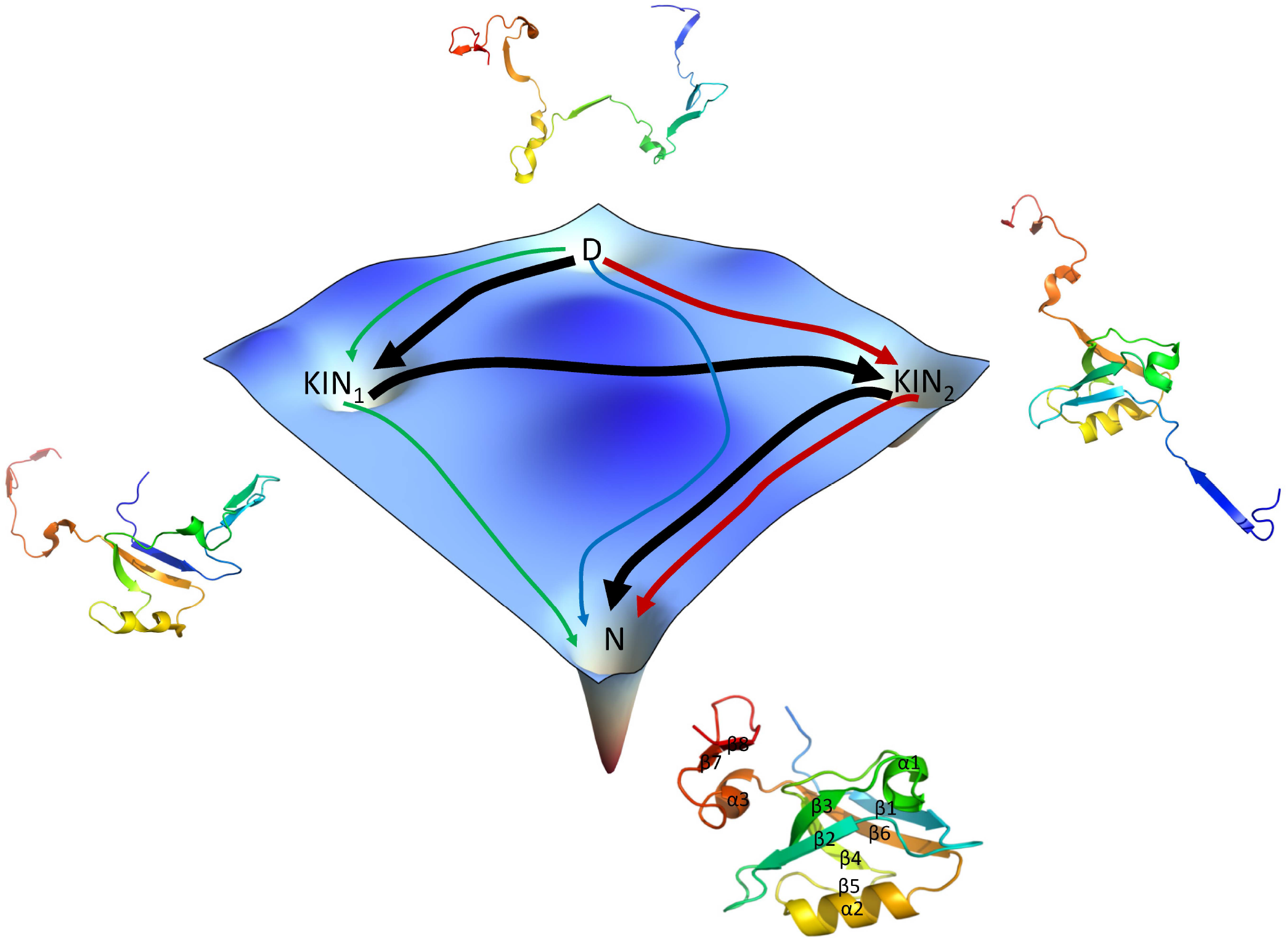
TOC Graphic

In contrast, the correlation between the calculated *f_Q_* between *I_EQ_* and *KIN*_2_ is high (Fig-8B). with the correlation coefficient, R=0.98. A linear fit of the line in Fig-8B gives *y* = *A*+*Bx* with *A* = 0.008±0.007, *B* = 1.175±0.021. Since the intercept, *A*, is close to zero, and the slope, *B*, is close to unity, we conclude that *KIN*_2_ and *I_EQ_* are the same intermediate species. Fig-8C shows, for *KIN*_1_, *I_EQ_* and *KIN*_2_, the average fraction of native contacts at every residue *f_Q_s* by color, vividly demonstrating the similarity between *KIN*_2_ and *I_EQ_*. In addition, *KIN*_2_ and *I_EQ_* are very similar to the native PDZ3 structure except that the *β*_1_ strand and *α*_3_ helix are not structured if the *β*_7_ and *β*_8_ strands are not included. These results show that in the later stages of folding the equilibrium and kinetic intermediates coincide, which is expected to hold for most, if not all, foldable proteins. In the later stages of folding, which corresponds to the stage after chain compaction, native-like interactions dominate, which was first shown using lattice models^45,46^ for which precise computations can be performed. Because native-like structures, which have considerable Boltzmann weight, dominate it follows that the structural features of *KIN*_2_ and *I_EQ_* should coincide.

Despite the simplicity of theory-based approach used here it is worth emphasizing again the validity of the SOP model from the following perspectives. First, the SOP model was not parameterized but is transferable, as we have demonstrated in a number of applications (see for example^9,12,43^). Second, the SOP model predictions for the thermodynamics are in excellent agreement with experiments not only for this protein but also for about ten proteins for which detailed comparisons have been made. Third, previous simulations using lattice models^45,46^ show that after the polypeptide chain collapses the transition to the native state is dominated by native-like interactions, which is consistent with our finding that the structural features of *KIN*_2_ and *I_EQ_* coincide and thus justifies the SOP model. For these reasons we believe that the major prediction that the fluxes through the pathways can be altered by changing the denaturant concentration is valid, and certainly amenable to experimental test as was done for adenylate kinase. ^30^

### Denaturants alter the connectivity between the metastable intermediates

The most interesting finding that denaturants alter the connectivity between the intermediates, and hence the fluxes through the distinct pathways, was already demonstrated in a most beautiful single-molecule fluorescence energy transfer (smFRET) experiment.^30^ Using adenylate kinase (ADK), a 214-residue two (or three) domain protein, and by collecting a large number of trajectories Haran and coworkers showed that during the folding process six metastable intermediates are populated. Most importantly, they showed using Hidden Markov Model analysis of the smFRET trajectories that the pathways traversed by ADK depend on the concentration of GdmCl. Both sequential (state i is connected to its neighbor *i*± 1) as well as non-sequential transitions, leading to parallel folding routes, are found during the folding process. The findings for ADK in experiments are qualitatively reflected in the simulated folding pathways of the smaller single domain PDZ domain. In particular, at the three concentrations of GdmCl we find both sequential and non-sequential connectivities. Just like the ADK study, we also find that a minor population of unfolded states directly reach the NBA, as predicted by the kinetic partitioning mechanism (KPM).^47^ Surprisingly, our simulations for PDZ show that KPM occurs only at 0.5M and 1.0M GdmCl but not in the absence of the denaturant. Overall, our simulations provide support to the discovery by Haran and coworkers ^30^ that fluxes through parallel folding pathways could be altered by changing the denaturant concentration.

It is interesting to compare this key finding^30^ to the reports of parallel unfolding routes in I27 induced by denaturants^48^ and SH3 domain by mechanical force (*F*).^49^ In both these studies the [*C*] or *F* dependence of ln *k_u_* (*k_u_* is the unfolding rate) exhibited upward curvature. Recently, we showed using theory that as long as the perturbation of the protein is linear in the external field (GdmCl or *F*) then upward curvature in the [ln *k_u_*, [*C*] or *F*] plot implies parallel unfolding pathways.^50^ In light of these observations and the present study, it would be most instructive to use *F* to probe the folding and unfolding of the PDZ domain. Such experiments would clarify if denaturant-induced intermediates coincide with those found under tension, which should be the case if the perturbation is linear in [*C*] and *F*.

Single molecule pulling experiments would provide insights into the nature of the TSEs and the modulation of fluxes through distinct pathways. There are two possible scenarios for forced-unfolding of PDZ3 in the presence of denaturants. The calculated free energy profiles in Fig. 3 suggest two possible outcomes for the single molecule pulling experiments. In the first case, we expect that the free energy profile could be described by an effective one-dimensional reaction coordinate with an outer barrier dominating at low forces and an inner barrier becoming important at high forces. ^51,52^ This scenario could hold good for force-induced unfolding of PDZ3, which would be consistent with the energy landscape inferred from ensemble experiments. ^27^ In this case there would be a change in the transition state position from a large (small) value at (low) (high) force. The more interesting scenario is that the location of the transition state in terms of the molecular extension, conjugate to the applied force, is an increasing function of force just as found for the unfolding of the SH3 domain. ^49^ In this case we would predict based on the free energy profiles in Fig. 3 that TSE1 would be dominant at low *F* and TSE2 at high *F*. Distinguishing between the two scenarios awaits single molecule pulling experiments.

## Conclusions

Using PDZ3 as another case study we have showcased the power of the SOP-MTM simulations in capturing accurately the thermodynamics of folding in the presence of denaturants. Although the unfolding rate in the absence of denaturants deviates substantially from experiments, the predicted folding rate in water is in excellent agreement with experiments. The major finding is that PDZ3 folds by parallel pathways with the crucial prediction that fluxes through the major pathways depend on the denaturant concentration. Single-molecule fluorescence energy transfer experiments could be used to validate our predictions. The two scenarios for parallel folding pathways, which lead to different predictions for the variation in the position of the transition states with changes in the mechanical force, can be distinguished using single molecule pulling experiments. Finally, the present work shows that the most practical, reasonably accurate, and currently the only way of taking the effects of denaturants into account is by using the SOP-MTM simulations. The transferability of this method has been established through numerous applications.

## Acknowledgement

ZL acknowledges financial support from the National Natural Science Foundation of China (11104015,11675017,11735005) and the Fundamental Research Funds for the Central Universities (2012LYB08). DT is grateful to the National Science Foundation (CHE 16-36424), the National Institutes of Health (R01 GM089685), and the Collie-Welch Regents chair (F0019) for supporting this work.

## References

(1) Thirumalai, D.; Liu, Z.; O’Brien, E. P.; Reddy, G. Protein folding: from theory to practice. Curr. Opin. Struct. Biol. 2013, 23, 22–29.

(2) Wallqvist, A.; Covell, D. G.; Thirumalai, D. Hydrophobic Interactions in Aqueous Urea Solutions with Implications for the Mechanism of Protein Denaturation. J. Am. Chem. Soc. 1998, 120, 427–428.

(3) Bennion, B.; Daggett, V. The molecular basis for the chemical denaturation of proteins by urea. Proc. Natl. Acad. Sci. 2003, 100, 5142–5147.

(4) O’Brien, E.; Dima, R.; Brooks, B.; Thirumalai, D. Interactions between hydrophobic and ionic solutes in aqueous guanidinium chloride and urea solutions: lessons for protein denaturation mechanism. J. Am. Chem. Soc. 2007, 129, 7346–7353.

(5) Canchi, D. R.; Paschek, D.; Garcia, A. E. Equilibrium Study of Protein Denaturation by Urea. J. Am. Chem. Soc. 2010, 132, 2338–2344.

(6) Hua, L.; Zhou, R.; Thirumalai, D.; Berne, B. Urea denaturation by stronger dispersion interactions with proteins t han water implies a 2-stage unfolding. Proc. Natl. Acad. Sci. U. S. A. 2008, 105, 16928–16933.

(7) Reddy, G.; Liu, Z.; Thirumalai, D. Denaturant-dependent folding of GFP. Proc. Natl. Acad. Sci. U. S. A. 2012, 109, 17832–17838.

(8) O’Brien, E. P.; Ziv, G.; Haran, G.; Brooks, B. R.; Thirumalai, D. Effects of denaturants and osmolytes on proteins are accurately predicted by the molecular transfer model. Proc. Natl. Acad. Sci. USA 2008, 105, 13403–13408.

(9) Liu, Z. X.; Reddy, G.; O’Brien, E. P.; Thirumalai, D. Collapse kinetics and chevron plots from simulations of denaturant-dependent folding of globular proteins. Proc. Natl. Acad. Sci. USA 2011, 108, 7787–7792.

(10) Liu, Z. X.; Reddy, G.; Thirumalai, D. Theory of the Molecular Transfer Model for Proteins with Applications to the Folding of the src-SH3 Domain. J. Phys. Chem. B 2012, 116, 6707–6716.

(11) Maity, H.; Reddy, G. Folding of Protein L with Implications for Collapse in the Denatured State Ensemble. J. Am. Chem. Soc. 2016, 138, 2609–2616.

(12) Reddy, G.; Thirumalai, D. Collapse Precedes Folding in Denaturant-Dependent Assembly of Ubiquitin. J. Phys. Chem. B 2017, 121, 995–1009.

(13) Maity, H.; Reddy, G. Thermodynamics and Kinetics of Single-Chain Monellin Folding with Structural Insights into Specific Collapse in the Denatured State Ensemble. J. Mol. Biol. 2017, 429, xxx–xxx.

(14) Onuchic, J. N.; Wolynes, P. G. Theory of protein folding. Curr. Opin. Struct. Biol. 2004, 14, 70–75.

(15) Thirumalai, D.; O’Brien, E. P.; Morrison, G.; Hyeon, C. Theoretical Perspectives on Protein Folding. Annu. Rev. Biophys. 2010, 39, 159–183.

(16) Dill, K.; Chan, H. From Levinthal to pathways to funnels. Nat. Struct. Biol. 1997, 4, 10–19.

(17) Dill, K. A.; MacCallum, J. L. The Protein-Folding Problem, 50 Years On. Science 2012, 338, 1042–1046.

(18) Doyle, D. A.; Lee, A.; Lewis, J.; Kim, E.; Sheng, M.; MacKinnon, R. Crystal structures of a complexed and peptide-free membrane protein-binding domain: Molecular basis of peptide recognition by PDZ. Cell 1996, 85, 1067–1076.

(19) Harris, B. Z.; Lim, W. A. Mechanism and role of PDZ domains in signaling complex assembly. J. Cell Sci. 2001, 114, 3219–3231.

(20) Kim, E. J.; Sheng, M. PDZ domain proteins of synapses. Nat. Rev. Neurosci. 2004, 5, 771–781.

(21) Jemth, P.; Gianni, S. PDZ domains: Folding and binding. Biochemistry 2007, 46, 8701–8708.

(22) Feng, H. Q.; Vu, N. D.; Bai, Y. W. Detection of a hidden folding intermediate of the third domain of PDZ. J. Mol. Biol. 2005, 346, 345–353.

(23) Gianni, S.; Engstrom, A.; Larsson, M.; Calosci, N.; Malatesta, F.; Eklund, L.; Ngang, C. C.; Travaglini-Allocatelli, C.; Jemth, P. The kinetics of PDZ domain-ligand interactions and implications for the binding mechanism. J. Biol. Chem. 2005, 280, 34805–34812.

(24) Chi, C. N.; Gianni, S.; Calosci, N.; Travaglini-Allocatelli, C.; Engstrom, A.; Jemth, P. A conserved folding mechanism for PDZ domains. FEBS Lett. 2007, 581, 1109–1113.

(25) Calosci, N.; Chi, C. N.; Richter, B.; Camilloni, C.; Engstrom, A.; Eklund, L.; Travaglini-Allocatelli, C.; Gianni, S.; Vendruscolo, M.; Jemth, P. Comparison of successive transition states for folding reveals alternative early folding pathways of two homologous proteins. Proc. Natl. Acad. Sci. USA 2008, 105, 19241–19246.

(26) Murciano-Calles, J.; Cobos, E. S.; Mateo, P. L.; Camara-Artigas, A.; Martinez, J. C. An Oligomeric Equilibrium Intermediate as the Precursory Nucleus of Globular and Fibrillar Supramacromolecular Assemblies in a PDZ Domain. Biophys. J. 2010, 99, 263–272.

(27) Gautier, C.; Visconti, L.; Jemth, P.; Gianni, S. Addressing the role of the α-helical extension in the folding of the third PDZ domain from PSD-95. Sci. Rep. 2017, 7, 12593.

(28) Murciano-Calles, J.; Guell-Bosch, J.; Villegas, S.; Martinez, J. C. Common features in the unfolding and misfolding of PDZ domains and beyond: the modulatory effect of domain swapping and extraelements. Sci. Rep. 2016, 6, 19242.

(29) Otzen, D.; Oliveberg, M. Salt-induced detour through compact regions of the protein folding landscape. Proc. Natl. Acad. Sci. 1999, 96, 11746–11751.

(30) Pirchi, M.; Ziv, G.; Riven, I.; Cohen, S. S.; Zohar, N.; Barak, Y.; Haran, G. Singlemolecule fluorescence spectroscopy maps the folding landscape of a large protein. Nat. Commun. 2011, 2, 493.

(31) Hyeon, C.; Thirumalai, D. Capturing the essence of folding and functions of biomolecules using coarse-grained models. Nat. Commun. 2011, 2, 487.

(32) Whitford, P. C.; Sanbonmatsu, K. Y.; Onuchic, J. N. Biomolecular dynamics: order-disorder transitions and energy landscapes. Rep. Prog. Phys 2012, 75, 076601.

(33) Pincus, D. L.; Cho, S. S.; Hyeon, C.; Thirumalai, D. Minimal Models for Proteins and RNA: From Folding to Function. Progress in Molecular Biology and Translationanl Science 2008, 84, 203–250.

(34) Sugita, Y.; Okamoto, Y. Replica-exchange molecular dynamics method for protein folding. Chem. Phys. Lett. 1999, 314, 141–151.

(35) Zhou, R. H.; Berne, B. J.; Germain, R. The free energy landscape for beta hairpin folding in explicit water. Proc. Natl. Acad. Sci. USA 2001, 98, 14931–14936.

(36) Sanbonmatsu, K. Y.; Garcia, A. E. Structure of Met-enkephalin in explicit aqueous solution using replica exchange molecular dynamics. Proteins 2002, 46, 225–234.

(37) Honeycutt, J. D.; Thirumalai, D. The Nature of Folded States of Globular-Proteins. Biopolymers 1992, 32, 695–709.

(38) Veitshans, T.; Klimov, D.; Thirumalai, D. Protein folding kinetics: Timescales, pathways and energy landscapes in terms of sequence-dependent properties. Fold. Des. 1997, 2, 1–22.

(39) Ermak, D. L.; McCammon, J. A. Brownian dynamics with hydrodynamic interactions. J. Chem. Phys. 1978, 69, 1352–1369.

(40) Liu, Z.; Reddy, G.; Thirumalai, D. Folding PDZ2 Domain Using the Molecular Transfer Model. J. Phys. Chem. B 2016, 120, 8090–8101.

(41) Noe, F.; Schutte, C.; Vanden-Eijnden, E.; Reich, L.; Weikl, T. R. Constructing the equilibrium ensemble of folding pathways from short off-equilibrium simulations. Proc. Natl. Acad. Sci. USA 2009, 106, 19011–19016.

(42) Li, W. F.; Terakawa, T.; Wang, W.; Takada, S. Energy landscape and multiroute folding of topologically complex proteins adenylate kinase and 2ouf-knot. Proc. Natl. Acad. Sci. USA 2012, 109, 17789–17794.

(43) Reddy, G.; Liu, Z. X.; Thirumalai, D. Denaturant-dependent folding of GFP. Proc. Natl. Acad. Sci. USA 2012, 109, 17832–17838.

(44) Cardenas, A. E.; Elber, R. Kinetics of cytochrome C folding: Atomically detailed simulations. Proteins: Structure, Function, and Genetics 2003, 51, 245–257.

(45) Camacho, C.; Thirumalai, D. Modeling the role of disulfide bonds in protein folding: Entropic barriers and pathways. Proteins: Structure, Function, and Bioinformatics 1995, 22, 27–40.

(46) Klimov, D. K.; Thirumalai, D. Multiple protein folding nuclei and the transition state ensemble in two-state proteins. Proteins - Struct. Funct. Gene. 2001, 43, 465–475.

(47) Guo, Z.; Thirumalai, D. Kinetics of Protein Folding: Nucleation Mechanism, Time Scales, and Pathways. Biopolymers 1995, 36, 83–102.

(48) Wright, C.; Lindorff-Larsen, K.; Randles, L.; Clarke, J. Parallel protein-unfolding pathways revealed and mapped. Nat. Struct. Biol. 2003, 10, 658–662.

(49) Jagannathan, B.; Elms, P. J.; Bustamante, C.; Marqusee, S. Direct observation of a force-induced switch in the anisotropic mechanical unfolding pathway of a protein. Proc. Natl. Acad. Sci. 2012, 109, 17820–17825.

(50) Zhuravlev, P. I.; Hinczewski, M.; Chakrabarti, S.; Marqusee, S.; Thirumalai, D. Force-dependent switch in protein unfolding pathways and transition-state movements. Proc. Natl. Acad. Sci. 2016, 113, E715–E724.

(51) Merkel, R.; Nassoy, P.; Leung, A.; Ritchie, K.; Evans, E. Energy landscapes of receptor-ligand bonds explored with dynamic force spectroscopy. Nature 1999, 397, 50–53.

(52) Hyeon, C.; Thirumalai, D. Multiple barriers in forced rupture of protein complexes. J. Chem. Phys. 2012, 137, 055103.

